# British Version of the Iowa Test of Consonant Perception

**DOI:** 10.1101/2024.09.04.611204

**Authors:** Xiaoxuan Guo, Ester Benzaquén, Emma Holmes, Inyong Choi, Bob McMurray, Doris-Eva Bamiou, Joel I Berger, Timothy D Griffiths

**Affiliations:** Biosciences Institute, Newcastle University, Newcastle upon Tyne, NE2 4HH, United Kingdom; Department of Speech Hearing and Phonetic Sciences, UCL; Department of Communication Sciences and Disorders, University of Iowa, Iowa City; Department of Psychological and Brain Sciences, University of Iowa, Iowa City; UCL Ear Institute; Human Brain Research Laboratory, Department of Neurosurgery, University of Iowa Hospitals and Clinics

## Abstract

The Iowa Test of Consonant Perception (ITCP) is a single-word closed-set speech- in-noise test with well-balanced phonetic features that provides a reliable testing option for real-world listening. **Objectives.** The current study aimed to establish a UK version of the test (B-ITCP) based on the British received pronunciation. **Design.** We conducted a validity test with 46 participants using the B-ITCP test, a sentence-in- noise test, and audiogram. **Results.** The B-ITCP demonstrated excellent test-retest reliability, cross-talker validity, and good convergent validity, consistent with the US results. **Conclusions.** These findings suggest that B-ITCP is a reliable measure of speech-in-noise perception, to facilitate comparative or combined studies in USA and UK. All materials (application and scripts) to run or construct the B-ITCP and ITCP are freely available online.

## 1 Introduction

Listening to speech in a noisy environment (Speech in Noise, or SiN) is common in everyday life, but it can be challenging for people with hearing impairments.

Traditional audiometric assessments of peripheral auditory ability, such as the pure-tone audiogram (PTA), predict SiN (Moore et al., 2020; Besser et al., 2015; Bochner et al., 2015; George et al., 2007), but they do not capture a large proportion of variance (Merten et al., 2022; Trine & Monson, 2020; Holmes & Griffiths, 2019; Jansen et al., 2014; Vermiglio et al., 2012). Therefore, measures that better simulate real-life listening conditions have been developed to assess word-in-noise or sentence-in-noise perception (see Reynard et al., 2022; Sanchez et al., 2022 for reviews of comparing different verbal SiN measures). SiN measures are not only an thus important part of audiometric battery in clinics to diagnose hearing loss, determine amplification choices, verify hearing aid performance (Taylor, 2003), but are also widely used in research that investigates cochlear synaptopathy, sound segregation, selective attention, ageing and cognitive impairments as well as other auditory cognitive mechanisms (Porto et al., 2023; Ryan et al., 2023; Guo et al., 2022; Le Prell, 2019; Billings & Madsen, 2018).

Understanding speech-in-noise is not the only challenge people face. When speech is heard from an unfamiliar dialect or accent in a noisy environment, this can also disproportionately impact people’s speech processing. This problem not only influences non-native speakers but also native speakers who are unfamiliar with different dialects and accents. For example, adult speakers of the Southern Standard British English have been found to show slower processing speed when listening to Glaswegian English especially in adverse listening conditions (Adank et al., 2009).

Similarly, while (Bent et al., 2021) show a decrement in a variety of native accents in normal hearing young adults, this effect is more pronounced in children: even as adults did not offer British vs. American accented speech, children can still struggle.

Aside from the word recognition accuracy, other aspects of speech processing are affected by accent. For example, The LiSN-S has similar normative data for British and American children but the talker advantage measure requires a corrective factor (Murphy et al., 2019). Speech processing also takes more effort when people are confronted with a less familiar accent (Van Engen & Peelle, 2014), suggesting accented speech (to a given listener) may tap a somewhat distinct set of cognitive and perceptual mechanisms than non-accented speech. Research also showed that older adults might have different cognitive strategies to younger adults when processing accented speech modulated by cognitive flexibility and inhibitory control (Ingvalson et al., 2017).

This work highlights a potential problem with the implementation of hearing assessments both in research and clinics: practitioners are often limited by the materials available to them and these materials might not be suitable for the population that they test. This is the case in UK audiology practice. Parmar et al., (2022) reported that only 20.4% of public sectors give speech tests in UK audiology practice. This is partly due to limited clinical resources but also because of the lack of widespread availability of materials geared towards British English. A large number of commonly used speech tests for hearing impairments used in the UK are not available in British English or validated with the British populations. A recent survey of British Audiologists and ENT surgeons of current clinical practice for the evaluation of auditory processing disorder (Bernard et al., 2024) reported that the most commonly used speech-based screening tools for adults were QuickSIN (Killion et al., 2004) and LiSN-S (Cameron & Dillon, 2007), and for children was LiSN-S, both recorded in American or Australian English only. Psycholinguistic work suggests that listeners not only tend to do poorly when confronted with speech from accents they are not familiar with (Van Engen & Peelle, 2014; Bent & Holt, 2013), McLaughlin et al. (2018) found that people’s relative skill at processing SiN did not even correlate with their skill at processing accented speech. Consequently speech-based tests can easily misidentify hearing problems by using a uniform standard (Dawes, 2011; Dawes & Bishop, 2007). From the patient’s perspective, developing and validating a well-designed SiN test is also in line with patient identified research priorities in the UK: individuals and families diagnosed with auditory processing disorder report the need for diagnostic tests as one of 3 top priorities (Agrawal et al., 2021); UK patients with mild to moderate hearing loss place the need for “realistic tests” of everyday hearing and potential use of SiN tests for hearing aid rehabilitation within the 15 top research questions that need to be answered (*James Lind Alliance*, 2024).

From a clinical perspective, it is clearly beneficial to make hearing tests available in the appropriate accent. From a research perspective, moreover, if matched versions of the same tests across accents were available, this could bring unique research opportunities. Research on the effect of accent and SiN can be investigated simultaneously. Such studies have been carried out but with individually recorded target stimuli and often a generic babble noise across accents that does not provide effective masking. Matched tests could make this more effective and offer standard measures that can be applied across studies.

Matched tests across accents also present an opportunity for larger public health-oriented work, taking advantage of “natural experiments” to assess the efficacy of various remediation approaches. For example, different criteria for medical interventions, such as cochlear implantation, are used in the US and UK. Candidacy for a cochlear implant in the UK, based on NICE guidelines, requires hearing loss ≥ 80 dB HL at two or more commonly-measured frequencies (between 500-4000 Hz). Contrastingly, common guidelines in the USA permit implantation when open-set sentence recognition in the best-aided condition is <60%, regardless of the degree of hearing loss. Consequently, a comparison of outcomes of hearing loss across the two populations could help reveal the best implantations strategies. However, such a direct comparison is difficulty without equivalent materials across the two dialects. Therefore, a SiN test that allows for better controlled comparisons between the US and UK populations could be important as it can potentially guide both clinical practices and research.

What type of SiN test would be a valuable addition to the current array of tests? The current zeitgeist in the field is to move towards the most ecologically rich form of such tasks – widely argued to be an open-set sentence-in-noise task. In this type of task (exemplified by the Hearing in Noise Task (Nilsson et al., 1994) or the AzBio Sentences in Noise Task (Spahr et al., 2012)), listeners hear a sentence (usually with low semantic predictability) and repeat it back to the experimenter. This is widely argued to much more precisely capture the range of skills that listeners might need in real-world SiN situations. Such materials are widely available for sentences across multiple accents (WILDCAT Corpus (Van Engen et al., 2010)). However, some of these skills are not auditory or even perceptual – the seemingly simple task of repeating a sentence (even in quiet) is a complex *cognitive* skill requiring lexical access, word recognition, sentence processing, language production along with embedded skills like working memory (Klem et al., 2015). Supporting this, sentence repetition *in quiet* is often seen as one of the best predictors of Developmental Language Disorder (Wang et al., 2022) —that is sentence recall is seen as a language measure, not a perceptual measure.

Some of these skills may also be affected in people who have hearing loss. For example, language may decline with age even in normal hearing individuals (Colby & McMurray, 2023; Payne et al., 2014; Waters & Caplan, 2001), or may be disrupted in children developing language with a hearing loss (Tomblin et al., 2015; Dunn et al., 2014). Consequently, single word tasks—if the words are well balanced from across the phonological space, may serve a valuable role in controlling some of this non perceptual variability and contributing to the research and clinical resources.

Similarly, while open set responding (in which a participant repeats back what is heard) is commonly seen to be more difficult and hence more sensitive to subtle differences (e.g., Clopper et al., 2006), it also poses speech production demands that may be challenging for some populations. In contrast, a well-balanced closed-set task – in which the response options are carefully chosen to reflect specific phonological dimensions of interest may be able to overcome this, maintaining a reasonable degree of difficulty while allowing the assessor to target particular dimensions of interest more precisely.

The Iowa Test of Consonant Perception (ITCP) was recently developed to overcome these concerns (Geller et al., 2021). It is a single-word, closed-set task that has a good balance of phonetic contrasts (expressed in the response options for each word) which covers the entire phonetic range of the English language. The original test showed very good test-retest reliability, as well as validity based on comparisons with the CNC word recognition test (Lehiste & Peterson, 1959) and AzBio sentence recognition test (Spahr et al., 2012).

This study sought to develop a British version of the same test using British English speakers with mainstream Standard Southern British accent. This is the modern equivalent of ‘Received Pronunciation’, which is widely used in education and the media. The development of B-ITCP aimed at benefiting both clinical practice and research.

To this end, we created a British version of the ITCP (British-ITCP or B-ITCP) for UK English speakers and validated it under laboratory conditions. The B-ITCP leverages the careful work of Geller et al (2021) in identifying an optimal and representative set of items and their response options, and simply replaces the audio with appropriate British accented versions of each stimulus. We evaluated performance accuracy, the test-retest reliability and the cross-talker validity to assess the reliability of the test itself. We also assessed the correlation between the pure-tone audiogram (PTA) and B-ITCP, and the correlation between B-ITCP and a sentence-in-babble (SiB) measure for the convergent validity (Holmes & Griffiths, 2019).

The B-ITCP is free and openly available to the community in the form of a testing APP and scripts that can be easily modified. It establishes a phonetically balanced measure of word-in-noise perception that, along with the freely available US ITCP stimuli, will allow direct comparisons between UK and US cohorts using a similar measure, and could facilitate combined studies in the two regions.

## 2 Methods

### 2.1 Participants

Forty-six English native speakers born and educated in the UK were recruited for the experiment (30 females, 16 males). Participants were excluded if they had a history of auditory disorders, speech or language disorders, developmental or neurological disorders or were taking psychotropic drugs. The PTA averaged across 0.25∼8kHz (in the left and right ears) of the sample was 13.92 dB HL, and the standard deviation (SD) was 8.42 dB HL. The average age was 48.65 (SD = 12.18).

A power analysis demonstrated that a sample size of 33 was required to show good reliability based on intraclass correlation coefficient (ICC) of greater than 0.75 and power of 80%. This was performed with an online sample size calculator (Arifin, 2024).

### 2.2 Materials and Design

Recordings were made by two native English speakers (one male and one female) with mainstream Standard Southern British accent. There are many accents in the UK and the received pronunciation was chosen because it is experienced by the majority of the UK population that is exposed to radio and television, even if it is not characteristic of their region. This selection of a standard accent is similar to that implemented by Geller et al. (2021).

The word list of the original ITCP test was recorded for each speaker (120 word sets per speaker). These are consonant-vowel-consonant words such as “ball-fall-shawl- wall”. Recordings were made in a sound-proof booth using a large-diaphragm condenser microphone (Rode NT1-A) with a pop filter placed in front. These recordings were made in Audacity (version 3.1.3), with a sampling rate of 44.1 kHz and 16-bit resolution. For both talkers, as per Geller et al. (2021), words were spoken as clearly as possible, at least twice with the carrier “he said [word]” and twice without. This phrase was included to help ensure uniform prosody and rate.

Offline, all words were imported into Audacity, the “Clip Fix” function was applied with 95% threshold for clipping and amplitude reduction overall by 5 dB (to allow for restored peaks). Noise reduction was then applied to the entire recording based on the noise profile for a silent period (with 12 dB reduction, sensitivity set to 6.00 and frequency smoothing set to 3). Each word exemplar was marked for cropping at the zero crossing, exported as a .wav file and then scaled to the same RMS level in Praat (version 6.2.14; Boersma, 2001) before being re-exported as a final “cleaned” .wav file. The mean duration of the words used was 0.51 s (±0.086 s)

The noise was extracted from an 8-talker babble soundtrack with 4 male and 4 female voices that lasts for 15 s in total. Importantly, this babble contained British voices, rather than the 8-talker babble with US accents included in Geller et al. (2021). In doing so, we attempted to create a stimulus set that provided as close as possible a direct analogue of the previous study, but for a different audience (native British listeners). Segments of the babble noise were taken randomly as a masker for the target word, which were always played 1 second before the target sound and stopped at the offset of the target words. The babble noise was mixed with the target sound with a -2 dB signal-to-noise ratio (SNR).

### 2.3 Procedure

The validation testing of B-ITCP was based on two sessions (Session A, Session B). The order of the two sessions was random, subject to participant availability. The two sessions were typically separated by 10 weeks (median duration = 80 days, range = 5∼356 days). In both sessions, researchers carried out separate audiometry, B-ITCP and SiB tests. The trial order of the B-ITCP task was kept the same for both sessions for all participants. The test sessions both started with a pure-tone audiogram (0.25 kHz ∼ 8 kHz), after which participants were asked to sit in a sound-proof booth in front of an LCD display (Dell Inc.) and were tested the B-ITCP test and a SiB test.

The two sessions are identical except for the SiB test: Session A tested the longer version of the SiB test and Session B had the same SiB test but shortened by half (this turned out to be unreliable and was not used in this study). Auditory stimuli were presented using headphones (Sennheiser HD 380 Pro) connected to an external sound card (RME FireFace UC). All computer tasks were programmed in MATLAB (R2021a, Mathworks, Natick, MA, United States). Participants were instructed to use either the keyboard or the mouse during the task.

The B-ITCP task consisted of 120 trials in total (shortened by half compared to the original ITCP task), with three blocks separated by short self-paced breaks. The whole test typically took 15 minutes to finish. Figure 1 contains a schematic of the trial structure. Each trial was up to 2 seconds long with a 1 second inter-trial interval. Half of the target words were spoken by the female speaker, while the other half was spoken by the male speaker. The order of the words was randomised between participants, but the same words were always spoken by the same speakers.

**Figure 1.**
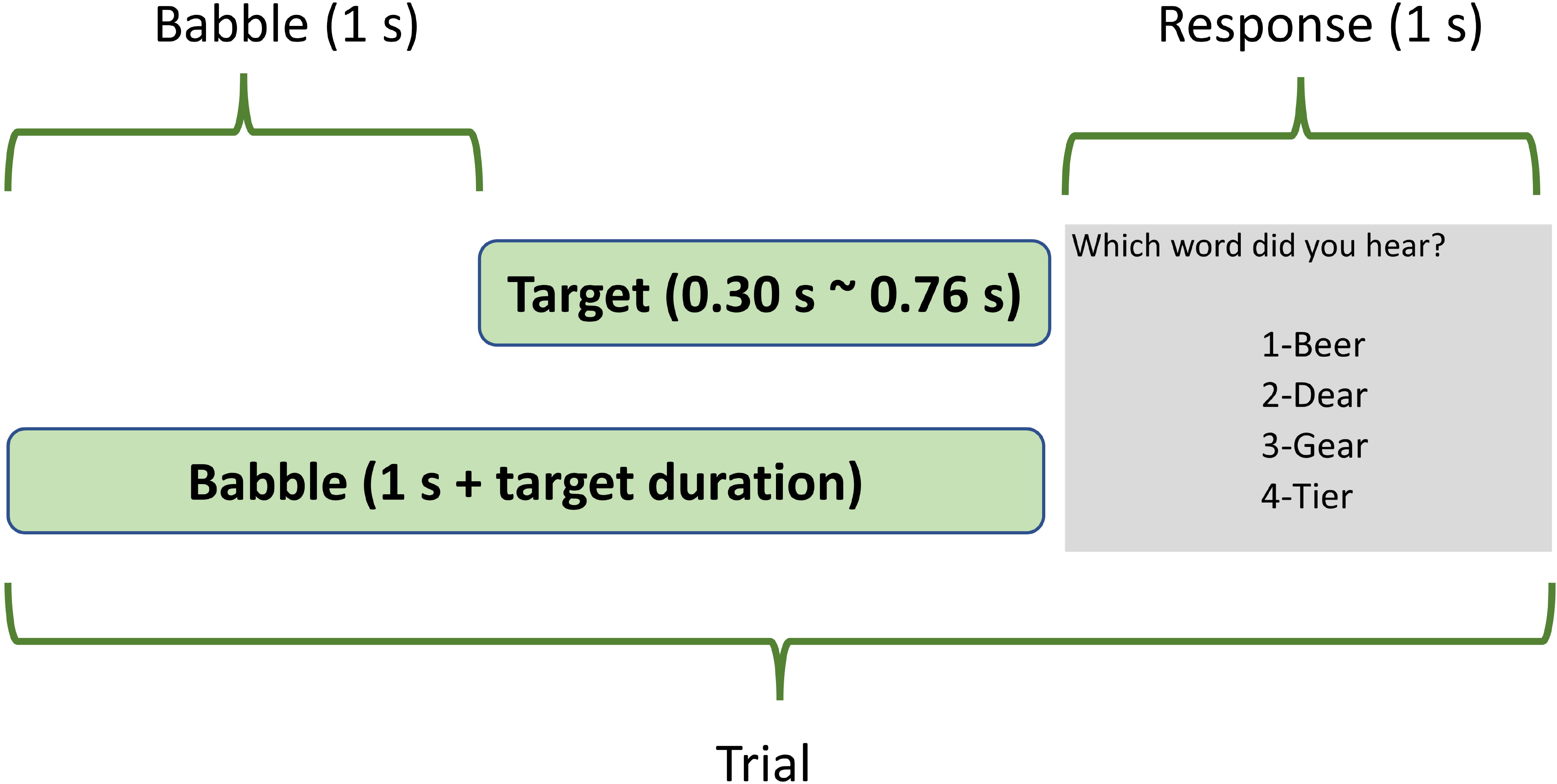
This figure shows an example of the trial structure of the B-ITCP test. The babble started 1s before the target, and the target typically lasted for 0.5s (range: 0.30 s to 0.76 s). After the target word was played, participants were shown 4 options on the screen and instructed to choose one of them based on what they just heard.

Participants were instructed to use the keyboard and choose the target word they heard out of four options on the screen that share the same vowel as the target word (e.g. “Beer-Dear-Gear-Tier”). The outcome measure used here for the B-ITCP was the proportion of words correctly identified (though the balanced structure of foils means the errors can be informative as well).

The sentence-in-babble (SiB) test was similar to that used by Holmes & Griffiths (2019). Target sentences were taken from the English version of the Oldenburg sentences and were recorded by a male speaker with mainstream received pronunciation of British English. Target sentences were structured as name-verb- number-adjective-noun; an example is “Alan brought four small desks”. The background noise was a 16-talker babble that had an onset 500ms before the target sentence. Participants were asked to repeat all five words from the target sentences: they were presented with a 5*10 matrix on the screen and were asked to select each of the five words from a list of 10 options with the mouse. They were required to guess if unsure. The test varied the SNR across trials. SNR started at 0 dB and proceeded with a one-down one-up staircase procedure depending on the participants response on each trial. SNR moved with a step size at 2 dB and but step size decreased to 0.5 dB after 3 reversals. The testing consisted of two interleaved runs, where each run had a different set of target sentences and terminated after 10 reversals. The median SNR of the last 6 reversals was taken for each run and both were averaged to compute participants thresholds.

### 2.4 Data Analysis

Data analysis was conducted in SPSS Statistics 29.0.1.0 and MATLAB R2021a. The results for both sessions were normally distributed, justifying the use of parametric tests. As the overall test design has been established with the previous validation study (Geller et al., 2021), the current study focused on test-retest reliability.

First, as the two sessions were not perfectly counterbalanced, we checked if there were learning effects or other outside influences that could lead to different performance on the two sessions. We compared the accuracy for each test between the two sessions with paired-sample t-tests.

Test-retest reliability was measured the same way as the ITCP validation (Geller et al., 2021), with the intraclass correlation coefficient (ICC), using a two-way random effects model (absolute agreement). ICC is commonly used to estimate the association between variable similar to Pearson correlation, but it considers both correlation and bias when assessing reproducibility (Liu et al., 2016). The absolute agreement measures are used to determine the level of agreement of raters, in this instance the scores of two B-ITCP testing (Koo & Li, 2016).

The relationship between B-ITCP and other speech and hearing measures was measured using Pearson’s correlations. Two pairs of correlations were assessed: PTA and B-ITCP (two sessions); B-ITCP and SiB (convergent validity check, for Session A only as the shorter SiB was not as reliable). A further cross-talker validity test was conducted by comparing the response to either the male or female speakers. A paired-sample t-test was used to assess if people responded differently to the two voices; the ICC further tests if the test can elicit reliable performance across talkers.

## 3 Results

The mean performance accuracy and standard deviations were extremely similar between the two sessions of B-ITCP: Mean (Session A) = 0.68 (SD = 0.08), Mean (Session B) = 0.67 (SD = 0.09). There was no significant difference in the mean performance between sessions: Mdiff = 0.005 (SD = 0.043), t (45) = 0.866, *p* = 0. 391. The SNR for SiB was Mean (Session A) M= -1.07 (SD = 1.44).

Figure 2 shows the correlation between PTA (averaged across 0.25 kHz to 8 kHz) and B-ITCP. Both sessions had large and significant negative correlations with a similar effect size: r (Session A) = -0.62 (p<0.001), r (Session B) = -0.56 (p<0.001). Note that the negative correlation is predicted since PTA is scaled such that a lower PTA indicates better hearing, while the B-ITCP is scaled such that higher scores indicates better performance.

**Figure 2.**
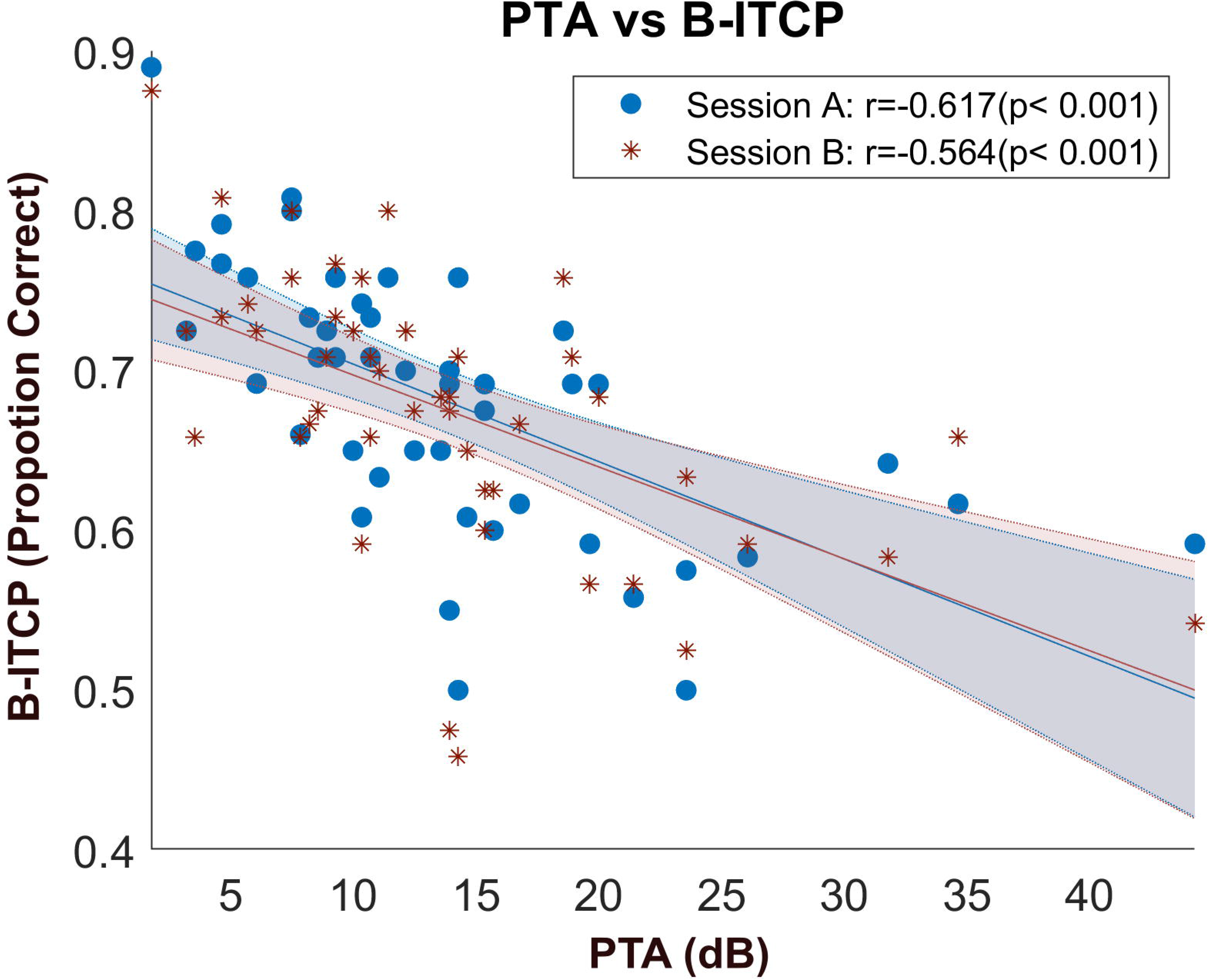
Scatterplot of PTA and B-ITCP of the two sessions. The correlation for Session A is in blue and for Session B is in red (the lines of best fit and error areas of the two sessions are in their respective colour as well). PTA results was taken from Session A. The x-axis plotted the PTA results in dB SPL, and the y-axis plotted B-ITCP results measured in the proportion of correct answers overall.

### 3.1 Test-Retest Reliability

We next examined the test-retest reliability of B-ITCP by calculating the ICC between the two sessions. The scatterplot (Figure 3) displays the close relationship between performance on the two sessions. This is further evidenced by the ICC results (Table 1) that showed excellent reliability of RICC = 0.93, which exceeds that of the original ITCP test-retest reliability of RICC = 0.80.

**Figure 3.**
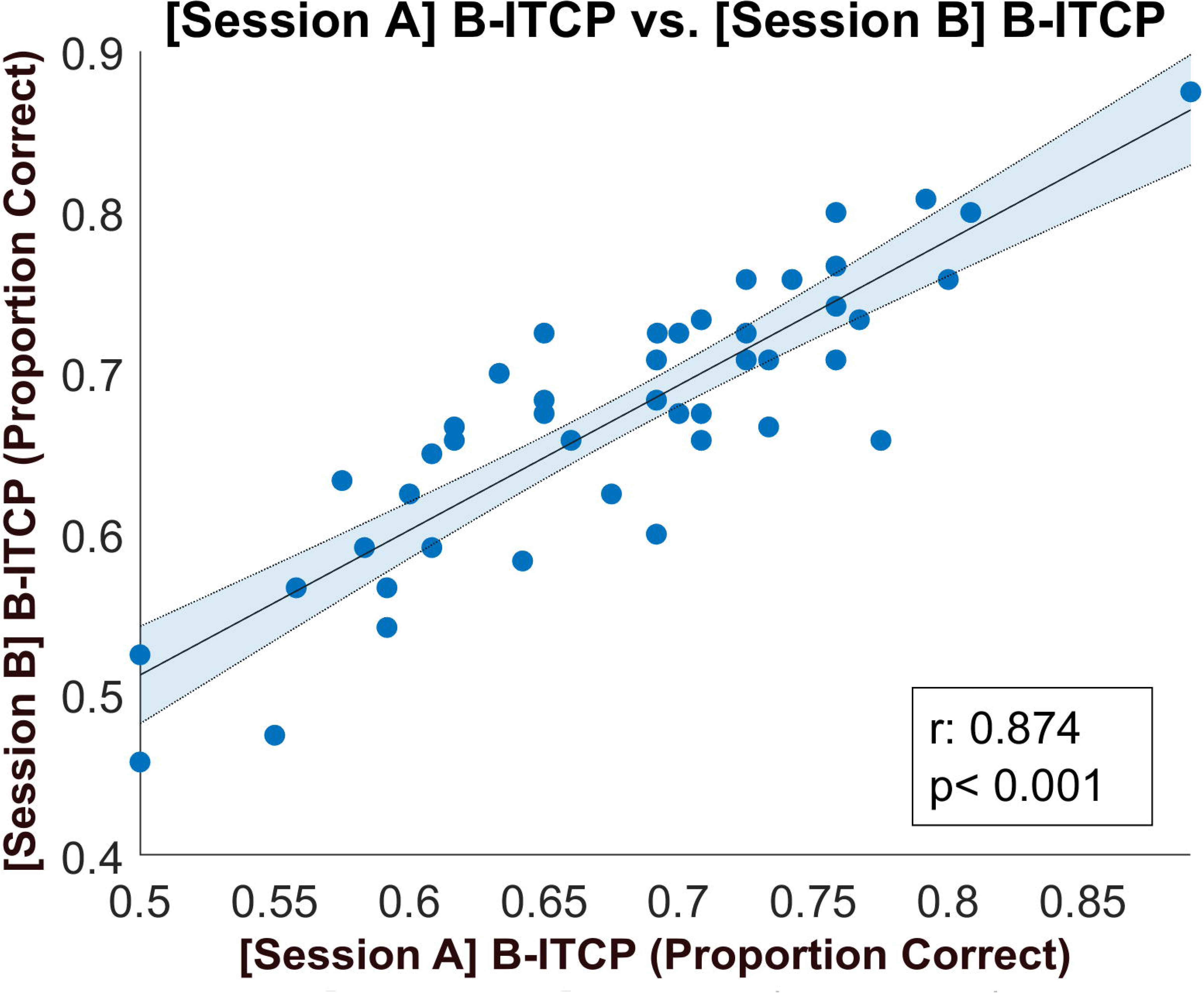
The scatterplot shows the association of the performance on B-ITCP in the two sessions. The x-axis represents the scores obtained from Session A and the y-axis represents the scores from Session B. Pearson’s r and a p value for a bivariate correlation are shown on the plot as well. The line of best fit is plotted in black with the error area shaded in blue.

### 3.2 Cross-Talker Validity

The cross-talker validity test showed that responses on the two sessions to either the female or the male voice did not differ significantly (M (Female Talker) = 0.68, SD (Female Talker) = 0.07; M (Male Talker) = 0.67, SD (Male Talker) = 0.08; t (45) = 1.82, p = 0.075). ICC showed a good reliability score as well: RICC = 0.79, p < 0.001.

### 3.3 Convergent Validity

The correlation between B-ITCP and SiB was -0.76 (*p* < 0.001), see Figure 4 for details. As with the PTA, SiB is scaled as a threshold, so the negative correlation is predicted.

**Figure 4.**
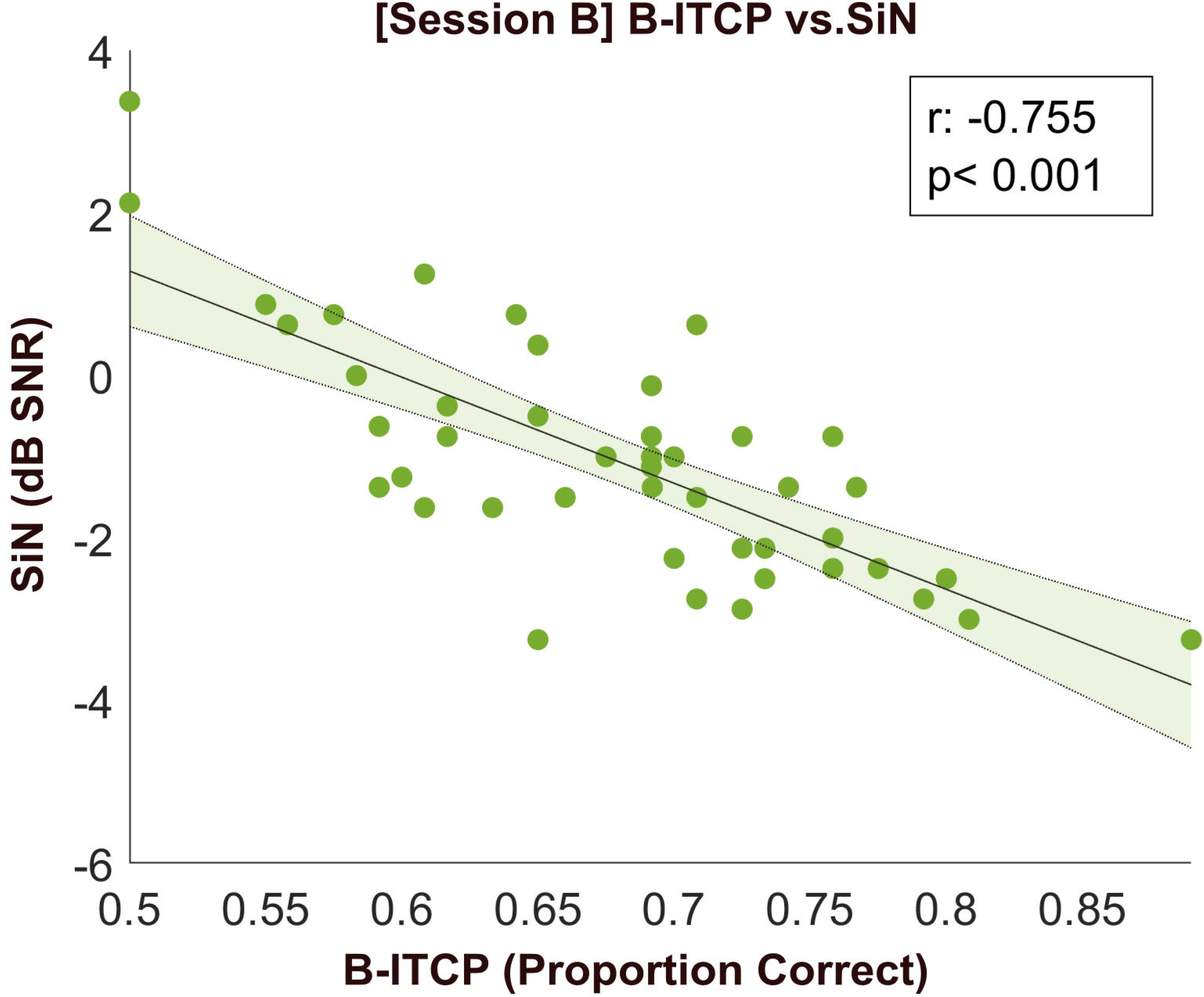
Scatterplot for bivariate correlations between B-ITCP and SiB. B-ITCP results are in proportion correct (x-axis) and SiB in dB SNR (y-axis). The line of best fit is plotted in black with the error area shaded in green.

## 4. **Discussion**

The performance data of B-ITCP showed a Gaussian distribution and achieved a reasonable level of accuracy (around 68% compared to 73% correct reported in Geller et al, 2021). Thus, the B-ITCP meets the minimal criteria for useful measure. While one of the goals of this study is to establish a test that can elicit comparable results from the UK and US, the performance accuracy of the current study cannot be directly compared with the ITCP results as the subject cohort and test parameters used here were not tightly matched with the ITCP study (which was validated online and tested all words in four subjects). To develop an equivalent test across the UK-US, further studies are needed which better align the detailed design of the study and the subject populations.

Further, the comparison of the mean accuracy between the two sessions showed no significant difference on performance of the two sessions. This means it is unlikely that the results were contaminated by learning or order effects. This is in part due to the unique design features of the ITCP closed-set task in which each of the four items that comprise a response set are used as the target (and they can be used multiple times across talkers). Consequently, subjects cannot learn which item is the correct response for a given set – they must process the stimulus.

We also demonstrated that PTA could predict B-ITCP performance in both testing sessions, which is consistent with our hypothesis and the literature discussed previously (Moore et al., 2020; Besser et al., 2015; Bochner et al., 2015; George et al., 2007).

Both the bivariate correlation and the ICC outcome demonstrated excellent test-retest reliability (ICC = 0.93). This means that the B-ITCP test can obtain a representative and stable assessment of SiN ability over time, which allows for both cross-sectional or longitudinal studies. Again, the ICC score is consistent with the previous results from Iowa (ICC = 0.80), but higher test-retest reliability was obtained in this study. One potential explanation of the higher ICC score in this study is that the validation for B-ITCP took place in laboratory conditions, but the ITCP validation test was carried out online where audio presentation, background noise and distraction cannot be as well controlled. A comparison of online and lab testing carried out by Bridges and colleagues found that online testing for both visual and auditory modality tended to generate lower precision and more variability in performance (Bridges et al., 2020). The researchers argued that such discrepancy in results between the two modalities would not invalidate online auditory research, but it did mean that validating online results was necessary. As the current analysis relies heavily on performance stability, it is expected that a more controlled environment will lead to higher ICC score. However, the online ITCP still achieved a very good ICC score (0.80), suggesting that the test can be reliably used online as well as in the lab.

The cross-talker validity assessments were carried out to ensure that each talker was representative of the whole. This was important as to obtain a shorter test, half of the stimuli were presented in each voice this contrasts with the original ITCP where full list of words was heard in both female voices). The shortened version is good for time-limited testing in the clinics or as a part of a lengthy auditory experiment but raised a concern over potentially less balanced results as people might respond differently to the acoustic/phonetic features of the two voices.

However, the non-significant t-test showed that participants did not perform differently, suggesting that the shortened version can provide a reliable assessment of people’s SiN ability. Additionally, the high ICC score highlighted an important aspect of validity check for SiN tests — checking if the selected talkers are representative of all the talkers who speak the language. All speech-based auditory tests work around the assumption that people would respond to the recorded talkers the same way as they respond to the typical speakers of that language. The cross- talker ICC test showed that this cohort did give highly uniform responses to the two randomly chosen speakers. It is therefore reasonable to assume that B-ITCP is a generalisable SiN test and people’s performance on B-ITCP is representative of their real-life listening.

A further assessment of the validity of B-ITCP against the SiN measure found that B- ITCP correlated strongly with the Oldenburg sentence-in-noise measure. This finding is consistent with the ITCP study (Geller et al., 2021), which has established a strong association between ITCP and other standardised SiN tests based on sentences.

This consistency in the correlation of (B-) ITCP with other SiN measures in the two validation studies (US and UK) suggest that first, the results are less likely to be due to other non-specific effects such as motivation and arousal, which can cause the participants performing better on all tests. Second, the (B-) ITCP can give very similar clinical assessment results to patients’ real-world listening ability despite that sentence-level SiN measures are thought to be more ecological. The fact that this closed-set word-level SiN test is shorter and engages a ‘purer’ auditory speech segregation process also adds to the benefit of using the test when sentence-level tests pose a problem.

As highlighted earlier, the development of a comparable speech-in-noise test in the UK and USA would allow for comparisons between two countries with very different criteria for interventions. The B-ITCP and the ITCP potentially represent two tests that can serve this purpose. Studies implementing both of these tests in similar populations of native listeners and with similar testing conditions could provide evidence for appropriate adjustments of policies or criteria in either locale, dependent on optimizing potential benefits for individuals with hearing difficulties. This includes benefits that are not detected via commonly-used peripheral measures, such as cortical changes that are often somewhat independent of cochlear function and reflect higher-level processing deficits (Berger et al., 2023; Choi et al., 2023), which are relevant to various aspects of auditory scene analysis. However, the current experiment only assessed the reliability of the test. To establish age-scaled normative scores, further testing is needed on a wider population, including a wider range of age and hearing sensitivity.

In conclusion, this study shows that British-ITCP test has excellent reliability, convergent validity, and generalisability. The shortened version as used in this study provides a good solution for a quick clinical SiN assessment. The full version can be used for research across the UK and US for a more comprehensive test. Both versions are freely available on our OSF page, and researchers can tailor the test based on their preferences.

## Acknowledgments

This study was supported by the Medical Research Council. The funder was not involved in the study design, data collection, data analysis, or reporting.

## Notes

The MATLAB scripts for running or constructing your own versions of (B-) ITCP are freely available at https://osf.io/53jsg/ (Xiaoxuan Guo, Ester Benzaquén, Emma Holmes, Inyong Choi, Joel I Berger, Timothy D Griffiths. (2024). DOI 10.17605/OSF.IO/53JSG); the desktop application is available at https://osf.io/tqjuk/ (Benzaquén, E., Guo, X., & Griffiths, T. (2024, August 29). Iowa Test of Consonant Perception: Desktop app with US and British versions. https://doi.org/10.17605/OSF.IO/TQJUK).

## Financial disclosures/conflicts of interest

This work was supported by the MRC [grant number MR/T032553/1]. There are no conflicts of interest, financial, or otherwise.

The data that support the findings of this study are available upon request.

## Ethics

Experimental procedures were approved by the research ethics committee of Newcastle University and written informed consent was obtained from all participants. Ethics number: 10356/2018; principal investigator Timothy D Griffiths.

